# Multi-site temporal control of optogenetic stimulation enhances firing frequencies in peripheral nerves

**DOI:** 10.64898/2026.05.15.724667

**Authors:** Tarah A. Welton, Tyler Currie, Arjun Fontaine, John Caldwell, Richard F. Weir, Diego Restrepo, Emily A. Gibson

## Abstract

We find that multi-site temporal control of optogenetic photostimulation in peripheral nerves can enhance firing rates by overcoming the intrinsic limitation of opsin photophysics. The benefits of multi-site optogenetic stimulation were demonstrated with three approaches: (1) *in silico* modeling, (2) *ex vivo* in the sciatic nerve, and (3) *in vivo* in the vagus nerve. An *in silico* model of multi-site optogenetic stimulation was developed in two Hodgkin and Huxley type neuron models, that supported our hypothesis. The *ex vivo* sciatic nerve showed an increase in firing frequency that is physiologically relevant for functional control. The technique was then applied *in vivo* for optogenetic vagus nerve stimulation resulting in significant changes in heart rate compared with standard methods of single-site stimulation. Improving the control of optogenetically induced neural firing will have broad impacts for future developments in optical nerve interfaces and brain-machine interfaces.

## Introduction

Optogenetics is a powerful technique that provides unprecedented control of excitable cells by genetic targeting of light activated ion channels in specific cell types (1). Since its inception in 2005 (2, 3), optogenetics has been widely adopted in biomedical research, and it is considered a promising approach for clinical treatment due to its inherent specificity. In the peripheral nervous system (PNS), optogenetics is being explored as an alternative to electrical stimulation for a number of applications including motor recruitment (4, 5), treatment of pain (6, 7), inflammation (8), and cardiac disease (9, 10).

In recent years, the catalog of excitatory opsins has expanded substantially to include variants with larger photocurrents for enhanced sensitivity (11-13), faster kinetics to increase response timing (14-18), and redshifted spectra (16, 19, 20) for deeper tissue penetration and multiplexing (21-23). These new opsin variants have been utilized successfully in both cell culture and the central nervous system (CNS) (13, 20, 22, 23). Comparatively, few of these new excitatory opsins have been successfully integrated into the PNS. Several research groups have reported challenges with viral expression of modified opsin variants in the peripheral nerve and have suggested that the immune system may play a role (24-27).

To date, channelrhodopsin-2 (ChR2) is still one of the most widely-used (27) and efficient (25, 26) excitatory opsins for PNS neuromodulation, and it has been used to study sensory feedback (7, 28) and motor control (4, 5, 25, 29, 30). However, ChR2 is limited by the relatively slow dynamics of its photocycle (31). When exposed to continuous or high frequency light, ChR2 rapidly becomes desensitized and recovers slowly (15, 22) resulting in a transient peak of photocurrent that rapidly decays to a steady state value (31, 32) and inadequate spiking at stimulation frequencies above 30 Hz (15) to 40 Hz (14, 23). Therefore, it is important to explore alternate methods to improve optogenetic firing in the PNS.

In this work, we investigated whether multi-site, temporal control of optical stimulation could be used to improve the firing frequency of opsins expressed in peripheral nerves. We developed and demonstrated two-site, or “drumbeat,” optogenetic stimulation (**Fig. 1**) in three experimental contexts: 1) computational modeling of spiking with two-site stimulation in Hodgkin and Huxley type neuron models, 2) electrophysiology recordings of optically induced compound action potentials (CAPs) in *ex vivo* mouse sciatic nerves using two optical fibers to deliver light to separate regions along the length of the nerve, and 3) recording changes in heart rate from optogenetic vagus nerve stimulation (VNS) using two light emitting diodes (LEDs) in anesthetized mice. Together these experiments showcase the efficacy of this technique and its utility in different systems. This proof-of-concept demonstration shows that multi-site optical stimulation can be utilized as an alternative method of enhancing optogenetic firing in peripheral nerves, leading to future improvements in optical control for nerve interfaces.

**Figure 1.**
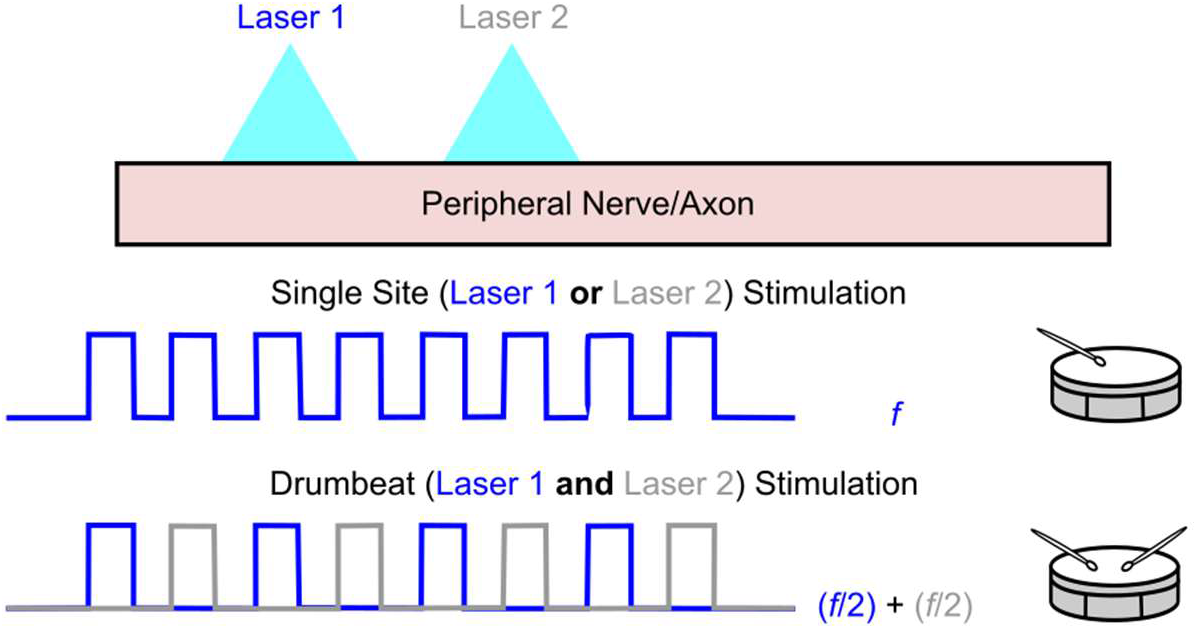
Schematic representation of two-site (“drumbeat”) optogenetic stimulation in a peripheral nerve. Single site stimulation uses only one site (either Laser 1 or Laser 2) at a specified frequency (*f*). Drumbeat stimulation uses two adjacent sites spaced along the nerve which are stimulated with separate lasers. Each site is stimulated at half the specified frequency (*f*/2) with Laser 2 temporally offset from Laser 1 by the period (1/*f*) so that their combined stimulation is *f*.

## Results

### Computational models of one and two-site optogenetic stimulation

Using the equations and modeling parameters from Stefanescu *et al*. (33), we evaluated single-site optogenetic stimulation, two-site “drumbeat” optogenetic stimulation, and pulsed electrical stimulation in single-compartment models of a fast-spiking interneuron and a hippocampal pyramidal neuron. The opsin photocycle was simulated using a four-state transition rate model (**Fig. 2a**) with two closed states (C_1_ and C_2_) and two open states (O_1_ and O_2_) (33, 34). In the absence of light, the total opsin population is in the dark-adapted, closed state (C_1_) (33). Once exposed to light, the population starts to transition into the other states at rates of P_1_, P_2_, G_r_, G_d1_, G_d2_, e_12_, and e_21_. O_1_ and O_2_ are the only conductive states where ionic current (I_ChR2_) is generated and is included with the other ionic currents in a Hodkin-Huxley type model (**Fig. 2b**). The magnitude of I_ChR2_ is described by **Equation 1**. V represents the membrane voltage, g_1_ and g_2_ represent the conductance of the O_1_ and O_2_ states respectively, ɣ = g_2_/g_1_, and O_1_ and O_2_ represent the fraction of the opsin population in the respective states. Since the O_1_ state is more highly conductive than the O_2_ state, ɣ is typically less than one (33, 34). The conductance of the O_1_ state (g_1_) represents the degree of ChR2 expression in a particular cell which is variable (33) and scales the amplitude of the resulting photocurrent.

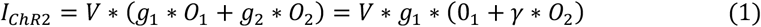

**Figure 2.**
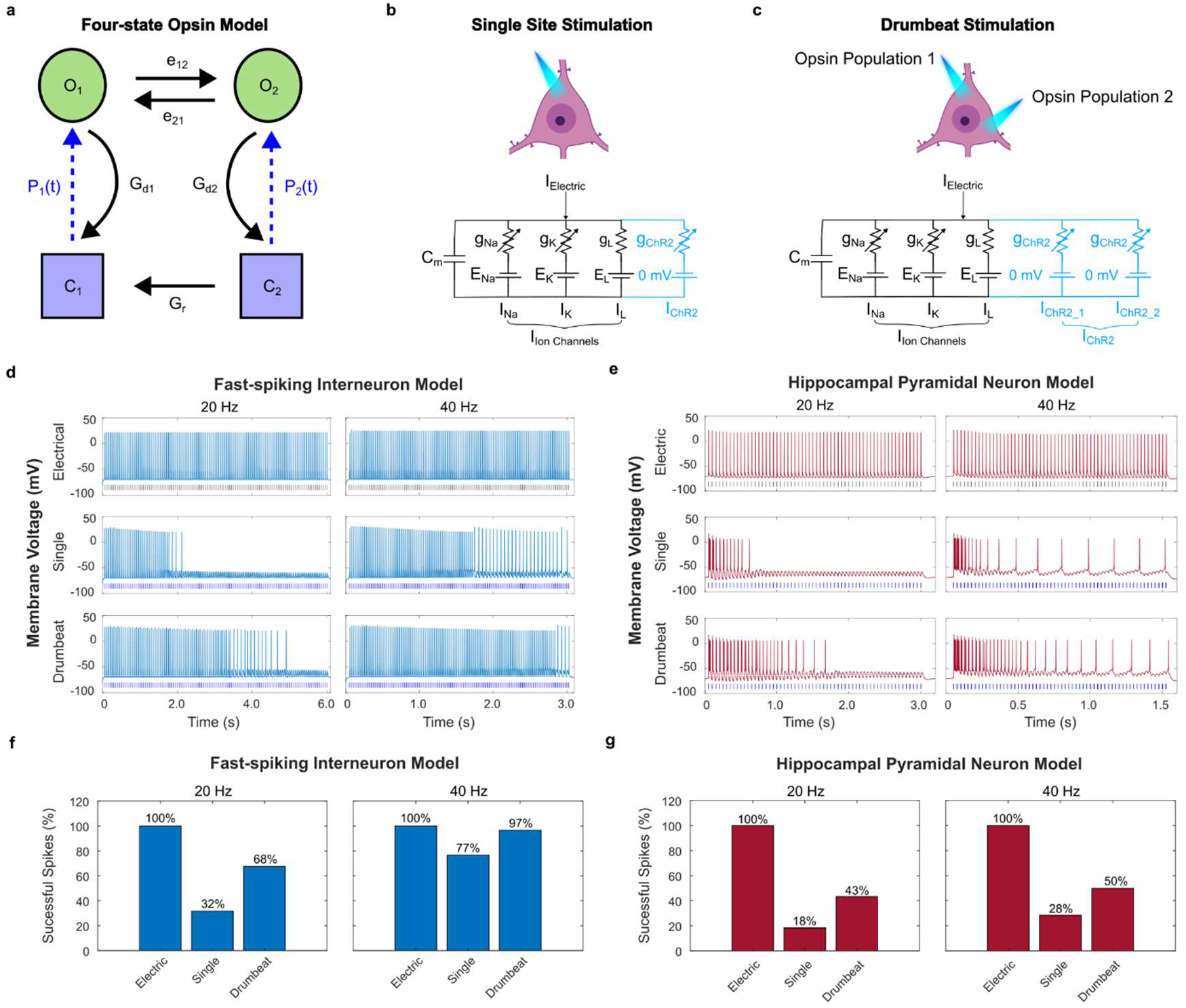
Computational modeling of single-site and drumbeat optogenetic stimulation in an interneuron and pyramidal cell. **(a)** Schematic of the four-state opsin model. C_1_, O_1_, O_2_, and C_2_ represent the fraction of opsin in each state at a given time. P_1_, P_2_, G_r_, G_d1_, G_d2_, e_12_, and e_21_ represent transition rates between the different states. Dotted lines represent light-induced transitions (P_1_ and P_2_). Adapted from Stefanescu *et al*. Fig. 1b(33). Representative schematic and circuit diagram analog of **(b)** single-site and **(c)** two-site drumbeat optogenetic stimulation in the Wang and Buzsaki fast-spiking interneuron. Cell schematics were created in BioRender. **(d)** Modeling results showing spiking in a fast-spiking interneuron at 20 and 40 Hz. The top panel shows the cell’s response to pulsed electrical stimulation (120 pulses, 0.2 ms pulse width, 72 µA/cm^2^). The middle panel shows spiking in response to single-site optogenetic stimulation of ChR2wt (120 pulses, 2 ms pulse width). The bottom panel shows optogenetic spiking in response to drumbeat stimulation of two populations of ChR2wt (each population gets 60 pulses, 2 ms pulse width). **(e)** Modeling ChR2wt spiking in a hippocampal pyramidal neuron at 20 and 40 Hz. The top panel shows the response to pulsed electrical stimulation (60 pulses, 0.2 ms pulse width, 15 µA/cm^2^). The middle panel shows optogenetic spiking to single site stimulation of ChR2wt (60 pulses, 2 ms pulse width). The bottom panel shows spiking in response to drumbeat stimulation of two populations of ChR2wt (each population gets 30 pulses, 2 ms pulse width). **(f**,**g)** Percent of successful spikes (%) in the interneuron and pyramidal neuron models respectively. Successful spikes are recorded as a single spike generated in response to one stimulation pulse.

Drumbeat optogenetic stimulation was implemented by adding a second, independent population of opsins to the model cell (**Fig. 2c**). This second population of opsins was modeled using the same parameters as the first population including the initial conditions, rate constants (P_1_, P_2_, G_r_, G_d1_, G_d2_, e_12_, e_21_), and opsin conductance values (g_1_, g_2_, ɣ). The two populations are independent because they are spatially separated, activated by different light sources, and the stimulation of the first population has no effect on the second population and vice versa. During drumbeat stimulation, both opsin populations were stimulated with the same light protocol (pulse width, frequency, and number of pulses), but the second opsin population was shifted in time by the period of the desired stimulation frequency. For example, drumbeat stimulation at 20 Hz required that both opsin populations were individually stimulated at frequencies of 10 Hz. The stimulation of the second opsin population was shifted in time by 0.05 seconds such that the combined stimulation of both populations stimulated the cell at 20 Hz (**Fig. 1**). The total photocurrent generated from drumbeat stimulation is described by **Equation 2**:

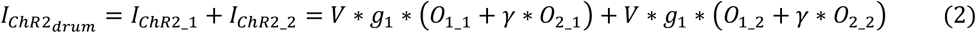

We validated the performance of our computational models by visually comparing our single-site stimulation modeling results to results from Stefanescu *et al*. (33) under the same stimulation conditions in both cell types (***SI Appendix* Fig. S1**). Next, we investigated the effect of drumbeat stimulation at 20 and 40 Hz. Compared to single-site stimulation, drumbeat stimulation visually improved spiking in the interneuron model (**Fig. 2d**) and the pyramidal neuron model (**Fig. 2e**) at both 20 and 40 Hz. The performance of each stimulation method was quantified by calculating the percentage of successful spikes where a successful spike was defined as one spike per delivered pulse. Extra spikes, as observed in the pyramidal cell model, were not counted towards the total. Out of 120 delivered pulses, the interneuron model successfully spiked 32% of the time at 20 Hz and 77% of the time at 40 Hz during single-site stimulation (**Fig. 2f**). Drumbeat stimulation increased successful spiking rates to 68% and 97% at 20 and 40 Hz respectively. Out of 60 delivered pulses, the pyramidal neuron model has a successfully spiking rate 18% of the time at 20 Hz and 28% of the time at 40 Hz during single-site stimulation. Drumbeat stimulation increased successful spiking to 43% and 50% at 20 and 40 Hz respectively (**Fig. 2g**). Surprisingly, both cell types produced more spikes at 40 Hz than they did at 20 Hz during single site and drumbeat stimulation. These results are likely because the membrane potential does not return to resting (-70 mV) between pulses at 40 Hz. This creates a slight plateau potential which temporarily increases the excitability of the cell resulting in increased spiking at higher frequencies. In summary, drumbeat stimulation effectively increased the number of successful optically-induced spikes in single-compartment computational models of a fast-spiking interneuron and a hippocampal pyramidal neuron.

### Experimental validation of multi-site optogenetic stimulation in ex vivo sciatic nerves

We experimentally tested single and two-site drumbeat optogenetic stimulation in *ex vivo* sciatic nerves. Sciatic nerves were excised from transgenic mice expressing ChR2 in cholinergic axons (ChAT-ChR2(H134R)-YFP). Custom-fabricated suction pipettes were secured to both ends of the nerve. The suction pipette on the sciatic-side of the nerve was used to deliver electrical stimulation while the suction pipette on the tibial-side was used to record compound action potentials (CAPs). Optical stimulation was delivered by pressing two optical cannulas against the side of the nerve using micromanipulators (**Fig. 3a**). Each cannula was connected to a separate 473 nm laser triggered by a transistor-transistor logic (TTL) pulse (***SI Appendix* Fig. S2**).

**Figure 3.**
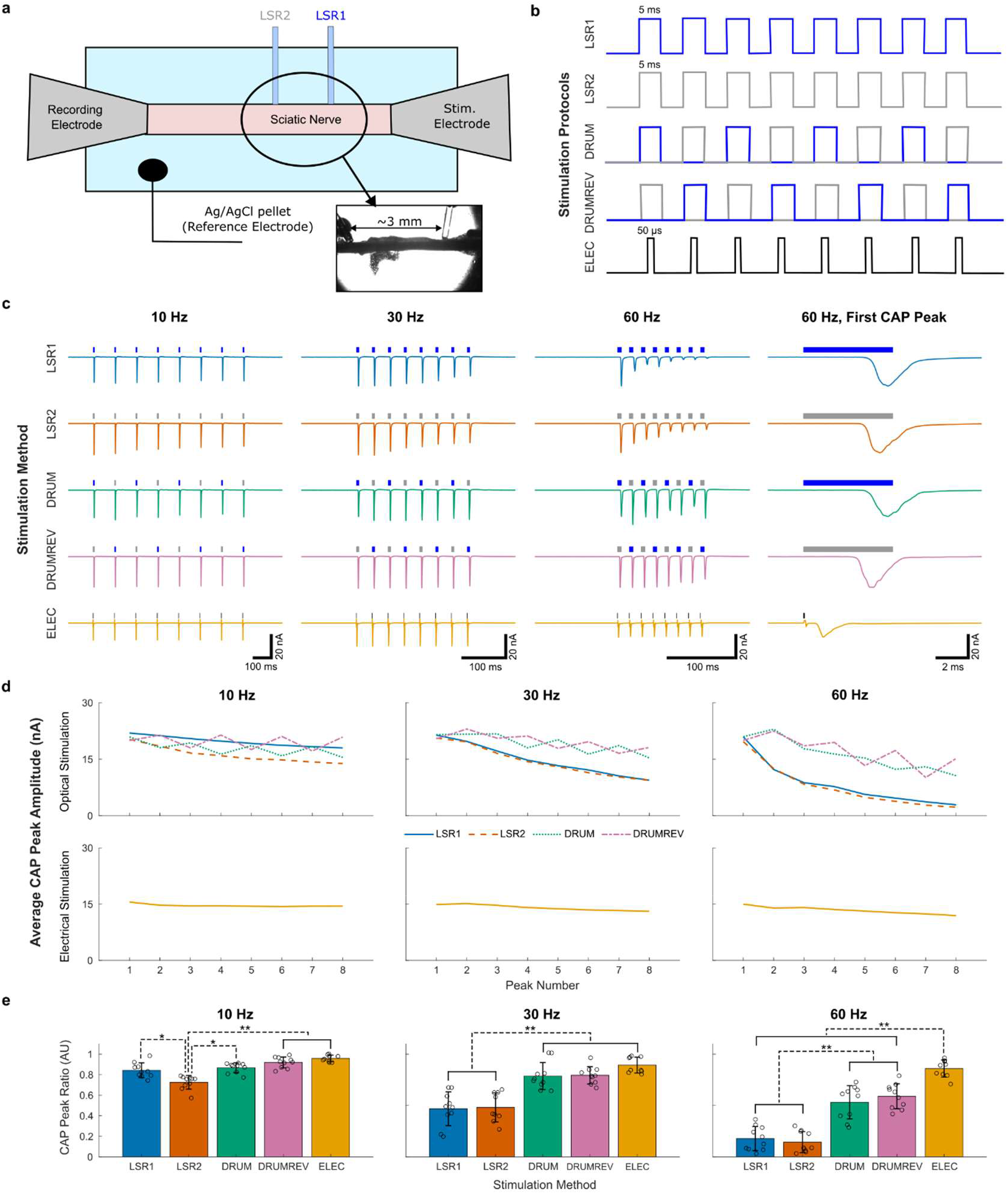
*Ex vivo* electrophysiology recordings of Compound Action Potentials (CAPs) from sciatic nerves. **(a)** Schematic of experimental set-up. Inset image shows a brightfield image acquired of the nerve and cannula placement. **(b)** Different stimulation time-sequence protocols. **(c)** Representative CAP traces from a single experiment in response to the different stimulation protocols. Each trace represents the average of five successive “sweeps” or trials to reduce noise. **(d)** The average CAP peak amplitude over all experimental trials plotted as function of peak number for optical stimulation methods (top panels) and electrical stimulation (bottom panels). **(e)** CAP peak amplitude ratios for all stimulation methods and frequencies. Bars represent means, error bars represent the standard deviation, circles represent individual data points, *p-values ≤ 0. 05, and **p-values ≤ 0.0001.

We tested three different stimulation frequencies (10, 30, and 60 Hz), five different stimulation methods (LSR1, LSR2, DRUM, DRUM REV, and ELEC) which are defined below, and two different inter-sweep intervals (20 and 60 seconds for optical methods and 2 and 10 seconds for electrical methods). For each stimulation frequency, we delivered a total of eight successive pulses with a width of 5 ms and 50 µs for optical and electrical modalities respectively. Stimulation was performed using: 1) laser one only (LSR1); 2) laser two only (LSR2); 3) drumbeat stimulation starting with laser one (DRUM); 4) reverse drumbeat stimulation starting with laser two (DRUM REV); 5) and electrical stimulation (ELEC) (**Fig. 3b**). The inter-sweep interval refers to the time delay between successive sweeps (or trials) of 8-pulse recordings. For every set of experimental conditions, we collected five, one-second-long sweeps which were averaged to reduce noise. For each experiment, we tested between 27 and 30 different experimental combinations for a single sciatic nerve over ∼2 hours. The experimental order was determined randomly prior to the experiment to account for potential order effects (35).

Representative CAP traces for each stimulation method and frequency from one experiment are shown in **Fig. 3c**. The optically induced CAP peaks show a reduced amplitude over eight successive pulses. As the stimulation frequency increases, the amplitude decay is more pronounced. For higher stimulation frequencies, drumbeat stimulation appeared to generate higher CAP peak amplitudes for subsequent pulses compared to single laser stimulation. Electrical stimulation produced very repeatable CAP peak amplitudes over eight successive pulses at all frequencies. However, the overall amplitude of the electrically induced CAPs varied depending on the recording order in the experimental sequence. For this particular experiment, the 30 Hz electrical trace was taken first and the intensity of the electrical stimulation was titrated to achieve a CAP peak amplitude of ∼ -18 nA (recording 8 of 27), the 10 Hz electrical trace was taken later in the experimental sequence (recording 20 of 27), and the 60 Hz recording was taken last (recording 26 of 27). Over the collection period that lasted approximately two hours, we observed that the overall amplitude of the electrically induced CAPs decreased with time.

To better compare the performance of the different stimulation methods over successive stimulation pulses, we plotted the CAP peak amplitude averaged over all experimental trials as a function of the stimulation pulse number (1 through 8) for each stimulation modality and frequency (**Fig. 3d**). Individual plots of all trials are shown in ***SI Appendix* Fig. S3a**. Single laser stimulation (LSR1 or LSR2) resulted in comparable CAP Peak Amplitude trends at 30 and 60 Hz. At 10 Hz, LSR2 appeared to underperform LSR1. At stimulation frequencies ≥ 30 Hz, drumbeat stimulation (DRUM and DRUM REV) elevated the CAP Peak Amplitudes and reduced the overall slope compared to single laser stimulation methods. However, drumbeat stimulation also introduced more peak-to-peak variability in CAP Peak Amplitude trends compared to single laser stimulation (***SI Appendix* Fig. S3a; *SI Appendix* Fig. S4a**) despite titrating the laser intensities at the start of the experiment (see Methods section). Electrical stimulation better maintained CAP peak amplitudes across successive peaks compared to all optical stimulation methods at all frequencies.

To quantify the performance of each stimulation method, we calculated the CAP Peak Ratio as the average amplitude of the last two CAP peaks (peaks 7 and 8) divided by the average amplitude of the first two CAP peaks (peaks 1 and 2). This metric was calculated using two peaks instead of one to better account for the peak-to-peak variability in the drumbeat traces. CAP Peak Ratios were compared using linear mixed-effects modeling where mouse number was treated as the random effect (see Statistics section). For data obtained under identical stimulation methods and frequencies, we found no statistically significant differences between the CAP Peak Ratios for the different inter-sweep intervals (p > 0.2 for optical stimulation methods and p > 0.3 for electrical stimulation) (***SI Appendix* Fig. S4**). Consequently, the inter-sweep interval was excluded as a factor of interest in subsequent analyses, and the data were pooled according to stimulation method and frequency.

CAP Peak Amplitude trends were assessed by evaluating the CAP Peak Ratio for the pooled data set (**Fig. 3e**). The CAP Peak Ratios elicited by drumbeat and electrical stimulation were significantly larger than those produced by single laser stimulation at frequencies ≥ 30 Hz (p ≤ 0.0001, n = 5 mice for optical stimulation methods and n = 6 for electrical stimulation). At 60 Hz, the electrical CAP Peak Ratio was significantly larger than those obtained with optical modalities (p ≤ 0.0001, n = 5 mice for optical stimulation methods and n = 6 mice for electrical stimulation). Maximal laser intensities used in *ex vivo* experiments (LSR1 = 62 mW/mm^2^ and LSR2 = 379 mW/mm^2^) did not elicit optical CAP responses in the absence of ChR2 expression (***SI Appendix* Fig. S3b**). In summary, drumbeat stimulation of *ex vivo* sciatic nerves resulted in significantly larger CAP peaks later in the pulse train compared to single laser stimulation at frequencies ≥ 30 Hz.

### *In vivo* two-site optogenetic vagus nerve stimulation (VNS)

*In vivo* experiments were performed acutely using custom 3D-printed nerve cuffs with an integrated light-emitting diode (LED) (**Fig. 4a**). ChAT-ChR2(H134R)-YFP mice (n = 3 mice) were anesthetized using isoflurane and placed on a heating pad (37.5°C) in a supine position. Electrical activity from the heart was recorded using electrocardiography (ECG). The left vagus nerve was surgically exposed (**Fig. 4b**) and LED1 was placed on the rostral portion of the vagus nerve using forceps. A preliminary three second, 30 Hz stimulation was performed to confirm that the cuff elicited a change in heart rate. Subsequently, LED2 was placed caudal to LED1 and tested for a confirmatory heart rate drop (**Fig. 4c**). After positioning, the light intensity of both LEDs was titrated to achieve a ∼10-20% drop in the mouse’s resting heart rate in response to three seconds of 30 Hz stimulation. This was defined as the “bradycardia threshold” or BCT (36).

**Figure 4.**
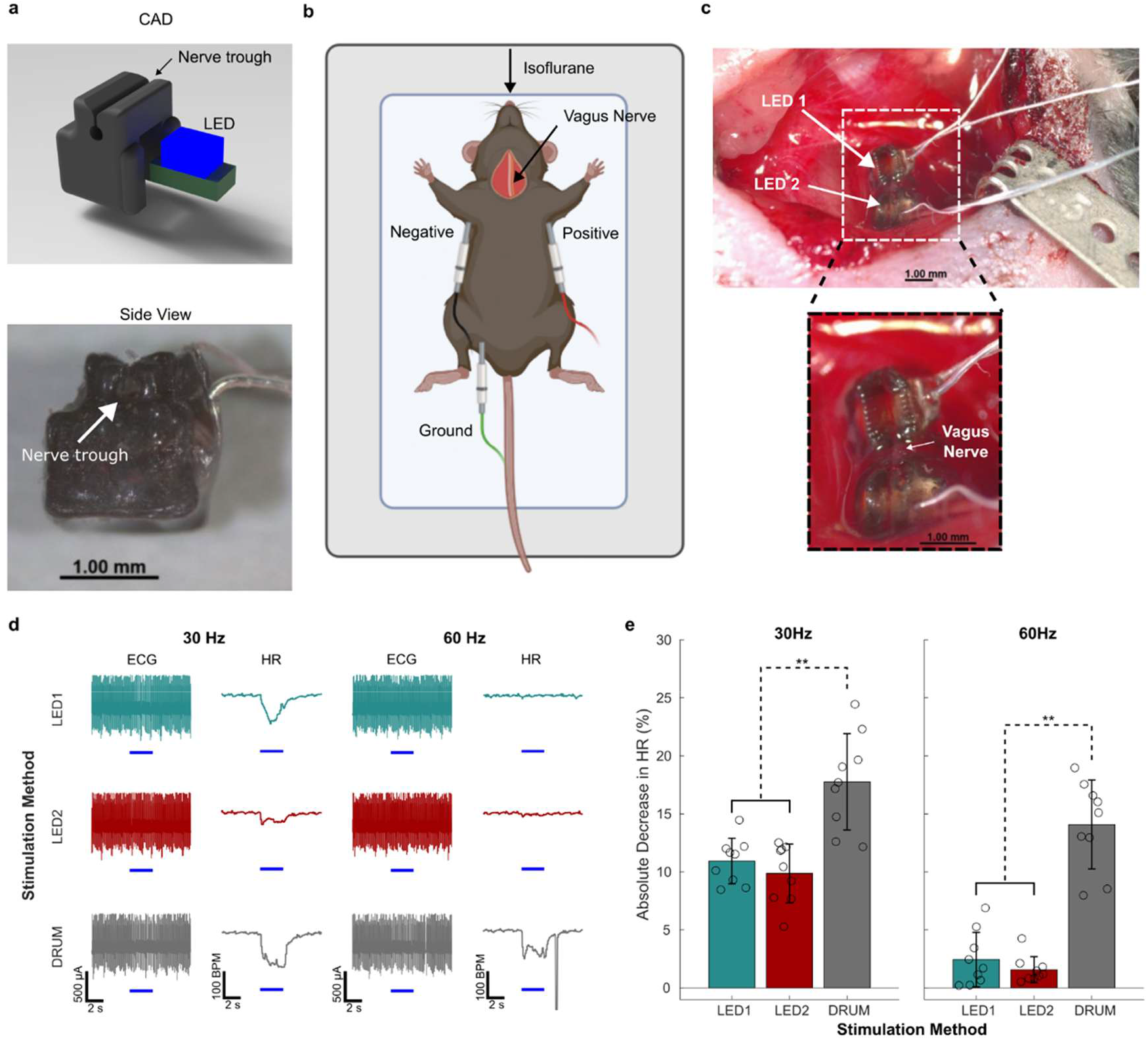
Acute *in vivo* drumbeat stimulation using custom-fabricated nerve cuffs. **(a)** CAD rendering of custom nerve cuffs and a photo of one of the fabricated nerve cuffs with the LED installed. **(b)** ECG electrode placement during *in vivo* recording. Mouse schematic was created in BioRender. **(c)** *In-situ* image of the nerve cuff placement and orientation. The inset picture shows a higher magnification image of the cuffs around the nerve. **(d)** Representative traces showing the raw ECG trace and the average heart rate (beats per minute) in response to three seconds of pulsed stimulation at different frequencies (5 ms pulses). **(e)** Absolute relative decrease in heart rate (%) for different stimulation methods and frequencies. Bars represent the means, error bars represent the standard deviation, circles represent individual data points, and ** p-values ≤ 0.0001.

Three trials were performed per animal. For each trial, we tested two stimulation frequencies (30 Hz and 60 Hz) and three optical stimulation methods: LED 1 only, LED 2 only, and drumbeat stimulation (DRUM: switching back and forth between LED1 and LED2, starting with LED1). This resulted in six stimulation combinations per trial. The experimental order was determined randomly prior to the experiment (36) with a two minute waiting period between successive stimulations to allow the mouse heart rate to fully recover. The BCT was reevaluated at the beginning of each trial, and the light intensity was adjusted if the heart rate dropped outside of the 10-20% BCT range. For all frequencies, stimulation was performed for a total of three seconds using 5 ms pulses.

Drumbeat stimulation resulted in larger decreases in heart rate compared to stimulation with a single LED at 30 and 60 Hz (**Fig. 4d**). This decrease was most pronounced at 60 Hz where stimulation with a single LED did not produce any visible change in heart rate. To quantify differences between stimulation methods, we calculated the relative change in heart rate during stimulation (ΔHR) (37) using **Equation 3**. HR_on_ represents the average heart rate during the three second stimulation window and HR_pre_ represents the average resting heart rate during the three second period just prior to stimulation.

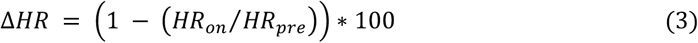

Calculating the relative change in heart rate accounts for differences in baseline heart rate between mice and multiple experimental timepoints (36) and the potential influence of different levels of isoflurane anesthesia on heart rate and residual vagal activity (38, 39) (**Fig. 4e**).

Each of the mice underwent three trials, and the results were analyzed using a linear mixed-effects model that treated mouse number as the random effect (see Statistics section). At 30 and 60 Hz, drumbeat stimulation resulted in significantly larger ΔHRs compared to single LED stimulation (p < 0.0001, n = 3 mice, 9 trials). Maximal LED intensities did not produce changes in heart rate in a control animal at either frequency (***SI Appendix* Fig. S5**).

## Discussion

This work demonstrates a novel method for optogenetic stimulation in peripheral nerves using two-site “drumbeat” stimulation. The efficacy of drumbeat stimulation was evaluated in *ex vivo* scatic nerves and *in vivo* vagus nerve stimulation (VNS) using an animal model with ChR2 expressed in cholinergic neurons. ChR2 is currently one of the most common and effective excitatory opsin variants used in the PNS (25, 27); however, its response is limited by its slow photokinetics, particularly at firing frequencies above 30 or 40 Hz. To get around this limitation, we hypothesized that drumbeat stimulation of multiple sites along a nerve would allow for more robust neuronal firing relative to single site optogenetic stimulation.

To explore this, we first implemented drumbeat stimulation by modifying two neuron-specific computational models of optogenetic stimulation to include two, independent opsin populations. In both the fast-spiking interneuron model and the hippocampal pyramidal neuron model, drumbeat stimulation of wildtype ChR2 increased spiking compared to single site stimulation at 20 and 40 Hz. Comparative analysis of these two neuron models demonstrates that the intrinsic biophysical properties of the cell greatly influence the neural response, despite the use of the same opsin variant (***SI Appendix* Fig. S6**). In the interneuron model, the amount of opsin in the more conductive open state (O1) gradually decreased over the course of successive stimulation pulses and the amount of opsin in the less conductive, light-adapted open state (O2) gradually increased. Alternatively, in the pyramidal neuron model, the amount of opsin in both O1 and O2 peaked at the start of the stimulation and decayed to a steady state value. In both models, the amount of opsin in the dark-adapted closed state (C1) decreased over successive pulses while the amount of opsin in the light-adapted closed state (C2) increased. Overall, the decay and growth of the different opsin populations in the interneuron model was more gradual and sustained than in the pyramidal neuron which allowed for successful action potential responses to twice as many stimulation pulses. Drumbeat stimulation effectively slowed the transition of opsins into their less responsive states (O2 and C2) by stimulating two populations at half the frequency of a single population. These initial results supported our hypothesis; however, single-compartment models are simple representations of neural activity and do not account for the spatial distribution of the cell and/or axon which could be an important variable for multi-site stimulation.

Experimentally, we tested the efficacy of drumbeat stimulation by optically stimulating two regions along the length of an *ex vivo* sciatic nerve from a ChAT-ChR2(H134R) mouse. These mice express the H134R variant of ChR2 in cholinergic neurons. Compared to wildtype ChR2, ChR2(H134R) has a slight reduction in desensitization and increased light sensitivity which produces larger photocurrents, but the channels also close at slower rates which makes it less temporally precise (22). We found that drumbeat stimulation produced more sustained CAP peak amplitudes compared to single site stimulation at frequencies ≥ 30 Hz. To our knowledge, the only other study that has investigated the frequency response of multi-site optogenetic stimulation in peripheral nerve is Moravec and Williams (40). They performed two and four-site stimulation of the peroneal nerve in anesthetized Thy1-ChR2(H134R) mice and recorded the electromyography (EMG) response from the tibialis anterior muscle. They found that multi-site stimulation at 75 Hz reduced the decay of the EMG response over a two second stimulation period which corroborates our *ex vivo* results.

We demonstrated the *in vivo* utility of drumbeat stimulation using optogenetic stimulation of the vagus nerve in an anesthetized mouse. Heart rate reduction is a well-documented feature of both electrical VNS (36, 37) and optogenetic stimulation of cholinergic vagal neurons (9, 41). Drumbeat stimulation resulted in a significant decrease in heart rate compared to single-site stimulation at 30 and 60 Hz (**Fig. 4e**). During electrical VNS, the reduction in heart rate scales with stimulation frequency, at least up to 50 Hz where the effect of frequency on heart rate plateaus (36). Optogenetic stimulation of cholinergic axons in the vagus nerve also results in heart rate reductions that scale with frequency up to 20 Hz (9). Drumbeat stimulation outperformed single-site stimulation at 30 Hz and completely restored the heart rate response at 60 Hz. This demonstrates that multi-site optogenetic VNS can be used to enhance opsin performance in a physiological setting. Although we only demonstrated optogenetic VNS using two stimulation sites, additional improvements may be possible with more than two sites. For example, incorporating an LED micro-array into a nerve cuff or fiber would allow for the stimulation of many sites along the length of the nerve (42, 43).

This proof-of-concept demonstration shows that multi-site temporally modulated optical stimulation can be utilized as a novel method of enhancing optogenetic firing in peripheral nerves, leading to future improvements in optical control for nerve interfaces. The method will allow researchers to perform optogenetic stimulation at physiological and clinically relevant frequencies. This work can be further expanded to include more than two stimulation sites, increasing the efficacy of optogenetic stimulation, approaching the temporal precision of electrical stimulation but with the significant benefit of genetic targeting to prevent the off-target effects of VNS.

## Materials and Methods

All experiments involving animals were conducted according to the NIH guidelines for animal research and were approved by the Institutional Animal Care and Use Committee at the University of Colorado Anschutz Medical Campus. Detailed materials and methods are available in SI Appendix.

## Data Availability

All data and code supporting this study will be made publicly available through GitHub after publication.

## Acknowledgments

The authors acknowledge Samuel Littich for conceptualizing the drumbeat stimulation paradigm and Katie Boncella for managing our animal colonies. We also acknowledge Dr. Cristin Welle for helpful feedback on experimental methods.

## Author Contributions

E.A.G. and D.R. conceptualized the project; J.C., R.F.W., A.F., D.R., and E.A.G. supervised the research; T.A.W. and T.C. performed the research. T.A.W analyzed the data and wrote the manuscript draft. All authors reviewed and edited the manuscript.

