## Supplemental Information for "Multi-site temporal control of optogenetic stimulation enhances firing frequencies in peripheral nerves"

### Supporting Information for
#### Multi-site temporal control of optogenetic stimulation enhances firing frequencies in peripheral nerves

Tarah A. Welton, Tyler Currie, Arjun Fontaine, John Caldwell, Richard F. Weir, Diego Restrepo, and Emily A. Gibson

Tarah Welton and Emily Gibson


**Supplementary Materials and Methods**

**Computational Modeling.** The computational models used in this paper were developed using the equations and modeling parameters from Stefanescu *et al.* (1). Stefanescu *et al.* integrated the three (2, 3) and four (4) state rate models of channelrhodopsin activity with two existing Hodgkin and Huxley type cell models of neuronal excitability: 1) the Wang and Buzsaki (1996) fast-spiking interneuron model (5) and 2) the Golomb *et al.* (2006) pyramidal hippocampal neuron model (6). Using experimental photocurrent data, they estimated the three and four state modeling parameters for multiple opsin variants (wildtype ChR2 (ChR2wt), ChETA (7), and ChRET/TC (8)) and qualitatively compared their modeling results with experimental cell firing data. They found that the four-state model better replicated experimental whole-cell patch clamp results compared to the three-state model. Stefanescu *et al.*'s MATLAB code is accessible through ModelDB (9) accension number 150804 (modeldb.science/150804).

Using MATLAB (version 2023a), we programmed single-site optogenetic stimulation in a fast-spiking interneuron and a pyramidal hippocampal neuron model. We expanded the Stefanescu *et al.* model to include: 1) more complex photostimulation protocols such as a tunable time delay at the start of the experiment and the option for multiple bouts of stimulation separated by a defined time interval, 2) two-site optogenetic (drumbeat) stimulation, and 3) pulsed electrical stimulation. In all optogenetic models, the four-state model of opsin kinetics (**Fig. 2a**) was used to simulate optogenetic activity. The rate constants and modeling parameters for the four-state opsin model are summarized in Table 3 of Stefanescu *et al.*, and the modeling parameters for the cells can be found in the text (1).

**Animals.** Adult ChAT-Cre (JAX# #006410) (10) x ChR2 mice (JAX# 024109) (11) between 17 and 30 weeks of age were used for all experiments. The *ex vivo* experimental cohort consisted of one female and four male mice that were heterozygous for both ChAT-Cre and ChR2 expression. A control experiment was performed in a male ChAT-Cre x ChR2 mouse that was wildtype for ChR2 expression. The *in vivo* experimental cohort consisted of two male and two female mice that were either heterozygous or homozygous for ChAT-Cre expression and homozygous for ChR2 expression. A control experiment was performed using a female VGlut2-Cre (JAX #028863) (12) x ChAT-Cre (JAX #006410) (10) x GCaMP6s (JAX # 024106) (13) mouse that was wildtype for VGlut2-Cre, ChAT-Cre, and GCaMP6s expression. Animals were housed in single-sex cages and provided with food and water *ad libitum*. Experimental protocols and animal care were performed in accordance with the Institutional Animal Care and Use Committee at the University of Colorado Anschutz Medical Campus.

**Suction Pipette Design.** Custom suction pipettes were hand fabricated by heating capillary glass (CAT 2-000-210, Drumond Scientific Company) over a Bunsen burner. The pipette fit was checked under a dissection scope. The stimulating electrode was composed of two uncoated silver wires (AG-15W, Warner Instruments). One of the wires ran through the center of the pipette while the other was wrapped around the outside (14). The recording electrode was a single silver wire extending through the center of the pipette. The wires were chlorinated by soaking them in bleach while simultaneously running a current through them using a 9V battery.

**Ex Vivo Nerve Electrophysiology.** Mice were euthanized using isoflurane anesthesia followed by cervical dislocation. The sciatic and tibial nerve branches were carefully dissected from the left and right hindlimbs and placed in room temperature, pH buffered mouse saline solution (126-mM NaCl, 5-mM KCl, 1.8-mM

$CaCl_2$, 1-mM $MgCl_2$, 10-mM MOPS Buffer at a pH of 7.2, and 30-mM Glucose) (14). The stock solution was titrated to a pH of 7.2-7.4 and an osmolarity of 300-310 mmol/kg (15).

A schematic of the *ex vivo* electrophysiology setup is depicted in ***SI Appendix*** **Fig. S2**. During recording, sciatic nerves were placed in a room temperature bath (~22°C, P6/PH6, Warner Instruments) of pH buffered mouse saline solution. Glass suction pipettes were used to deliver electrical stimulation to the sciatic-side of the nerve and record compound action potentials (CAPs) from the tibial-side of the nerve. An Ag/AgCl pellet (E205, Warner Instruments) was submerged in the bath to act as the reference electrode. Light pulses were delivered using two fiber optic cannulas (CFMC12L20, 0.39 NA, 200 µm core, Thorlabs). Cannulas were pressed flush against the nerve using two separate micromanipulators (MP-285, Sutter Instruments) at sites separated by ~3 mm. Each cannula was connected to a 473 nm laser (BL473T8-100FC, Shanghai Laser Optics Century Co. (SLOC)) via a patch cable (M81L01, 0.39 NA, 200 µm core, Thorlabs) using a quick-release interconnect (ADAF2, Thorlabs). After positioning the cannulas, the intensity of both lasers was titrated to achieve a CAP peak amplitude of ~ -18 nanoamperes (nA) for the first peak (LSR1: 19-62 $mW/mm^2$ and LSR2: 189-379 $mW/mm^2$). The intensity of the electrical stimulation was similarly titrated during the first electrical recording of the experiment (28-83 mV).

Experiments were performed inside a Faraday cage on an Axioskop 2 FS plus microscope (Zeiss). A long working distance air objective (A-Plan 2.5X/0.06, Zeiss) and camera (BFLY-U3-23S6M-C, FLIR) were used to confirm the cannula positions. A battery-operated stimulator (ISO-FLEX) delivered electrical stimulation via the suction electrode on the sciatic side of the nerve. CAP recordings were collected at 20 kHz using a CV-201A headstage, an Axon Digidata 1550A digitizer, an Axopatch 200A amplifier with a gain of 0.5 and a lowpass Bessel filter at 5 kHz, and Clampex Software (version 10.5.0.9, Molecular Devices). Due to the use of the CV-201A headstage, the maximum recordable current was limited to ±20 nA.

**3D Printed Nerve Cuffs.** The nerve cuffs were designed in SOLIDWORKS (version 2024, Dassault Systèmes) and 3D printed on the B9Core 530J printer (B9Creations) in BioRes-Silicone (B9R-BIO-SIL, B9Creations). The cuffs were cleaned with isopropyl alcohol and post-cured with 385-460nm light (B9Model Cure, B9Creations) in deionized water for ~10 minutes. The LEDs (C470DA2432-S3000-x, Cree) were sourced from eBay and arrived pre-soldered to a printed circuit board (PCB) board with wires. The original wires were removed and replaced with more flexible Teflon-coated wires (Sigmund Cohn Corp.) to reduce the strain on the cuffs *in situ*. The LED was inserted into the cuff from the bottom and secured with Loctite superglue. A secondary coat of Gorilla Glue Epoxy was added to better insulate the LED circuit board and soldered wire ends (**Fig. 4a**).

Before every experiment, the nerve cuffs were tested for electrical leaks by measuring the change in resistance between the air and a bath of 1x Phosphate Buffered Saline (PBS). If the resistance dropped when the cuff was submerged in saline, the cuff and wires were resealed with glue or a new cuff was used.

**LED Circuit Design.** The LEDs were controlled using a programmable Arduino board (Uno R3, Arduino) and custom code written using Arduino IDE (version 2.3.6, Arduino) software. The desired stimulation protocol (e.g. "LED1 at 30 Hz") was uploaded to the Arduino, and stimulation was manually triggered using a push button wired to a digital pin (as an input). Each of the LEDs were powered by their own digital pin (as an output) and were in series with a potentiometer (3590S-2-104L, Bourns) to control the light intensity (***SI Appendix*** **Fig. S7**).

**Electrocardiogram recordings.** A full schematic of the *in vivo* experimental setup can be found in ***SI Appendix*** **Fig. S7**. ECG data was amplified using a Bio Amp (FE231, AD Instruments), digitized with PowerLab 2/26 (PL2602, AD Instruments), and acquired using LabChart Pro (v8.1.30, AD Instruments). We used the second input channel on the digitizer to collect the TTL pulse train from the Arduino Uno. This allowed us to directly correlate changes in ECG and the time averaged heart rate data with stimulation.

**Data Analysis.** All data analysis was performed using MATLAB (version 2023a). Electrophysiology data was extracted from Axon Binary Files (ABF) using abfload.m(16). For each trace, the offset from zero was calculated by taking the average of the first 2000 data points before stimulation. The offset was subtracted from the overall trace. Five sweeps were performed per recording, and average CAP traces were calculated by taking the average of each of the five sweeps at each timepoint. From the average CAP traces, we calculated the CAP Peak Amplitude as the maximum value of each CAP peak. For electrical stimulation, we used linear interpolation (17) to remove the stimulation artifact prior to calculating the CAP Peak

Amplitude. From the CAP Peak Amplitude data, we calculated the CAP Peak Amplitude Ratio (the ratio of the last two peaks divided by the first two peaks).

*Ex vivo* drumbeat stimulation produced greater peak-to-peak variability in the CAP Peak Amplitude traces compared to single laser stimulation (**Fig. 3d; *SI Appendix* Fig. S3a**). This variability is likely the result of minor changes in the position of the optical cannulas relative to the nerve during the experiment due to micromanipulator drift and/or saline evaporation. When there was a significant drop in signal between one recording and the next, the position of the cannulas was adjusted and the light intensity was re-titrated if necessary. Since the bath was not perfused during experiments, evaporation caused the osmolarity of the bath to increase by ~110 mmol/kg over the course of a two-hour recording session. The effect of osmolarity on CAP traces was mitigated by the randomization procedure for each recording session. Furthermore, most of the drumbeat CAP peak variability (DRUM and DRUMREV) is associated with the operation of LSR2 (***SI Appendix* Fig. S3a**). The LSR2 cannula was positioned further down the nerve and away from the stimulating electrode which may have resulted in a positional shift over the course of the experiment (**Fig. 3a; *SI Appendix* Fig. S2**). LSR2 consistently required much higher laser intensities (189-379 $mW/mm^2$) than LSR1 (19-62 $mW/mm^2$) which is also indicative of suboptimal cannula positioning relative to the nerve.

ECG data was exported from LabChart Pro as a text file and opened in MATLAB for further analysis. Individual traces were identified using digital comments made during data collection, and the raw ECG signal and time averaged heart rate (beats per minute) were extracted for further analysis. We calculated the relative change in heart rate ($\Delta$HR) for each recording using **Equation 3**.

**Statistical Methods.** Statistical analysis was performed using RStudio (version R 4.4.1, R Core Team). Linear mixed-effects modes (LMMs) were used to analyze the *ex vivo* and *in vivo* data due to their ability to handle repeated measurements within subjects and unbalanced data sets (18, 19). LMMs were implemented using the lme4 package (20) with restricted maximum likelihood (REML) estimation. All LMM models were assessed by visually examining the Pearson residuals for normality and homoscedasticity using residual and Q-Q plots. Post-hoc, we used the emmeans package (21) to calculate the estimated marginal means, or the least-square means, of the LMMs. Bonferroni-adjusted pairwise comparisons were used to assess statistical differences between the stimulation methods at each frequency.

For the *ex vivo* data, the CAP Peak Ratio was the response variable, mouse was treated as the random effect, and frequency, stimulation method, and inter-sweep interval were specified as fixed effects. The *ex vivo* cohort consisted of six mice (five experimental and one control). Optical data were collected from all five experimental mice, except for a single missing data point for the LSR2, 30 Hz, 60 second inter-sweep interval condition. Electrical data were collected from all six mice. The 10 second electrical inter-sweep interval was collected for three of the six mice, and the 2 second electrical inter-sweep interval was collected from all six. To assess the effect of inter-sweep interval on the CAP Peak Ratio, we analyzed the optical and electrical data using separate models stratified by stimulation method and frequency (***SI Appendix* Fig. S4**). To assess the effect of each stimulation method at each frequency, the optical and electrical data were pooled into a single model (**Fig. 3d and 3e**). There were no major deviations in the normality or homoscedasticity of the models' residuals.

For the *in vivo* data, $\Delta$HR was the response variable, mouse was treated as the random effect, and frequency and stimulation method were specified as fixed effects. The *in vivo* cohort was comprised of four mice (three experimental and one control). Modeling mouse as a random effect resulted in a LMM with a singular fit with a random effect variance of zero. This is feasible since $\Delta$HR was normalized to the mouse's resting heart rate on a trial-by-trial basis. To confirm the modeling results, we refitted the model with mouse as a fixed effect instead of a random effect and got similar estimates and post-hoc inferences. In both cases, there were no major deviations in the normality or homoscedasticity of the models' residuals.

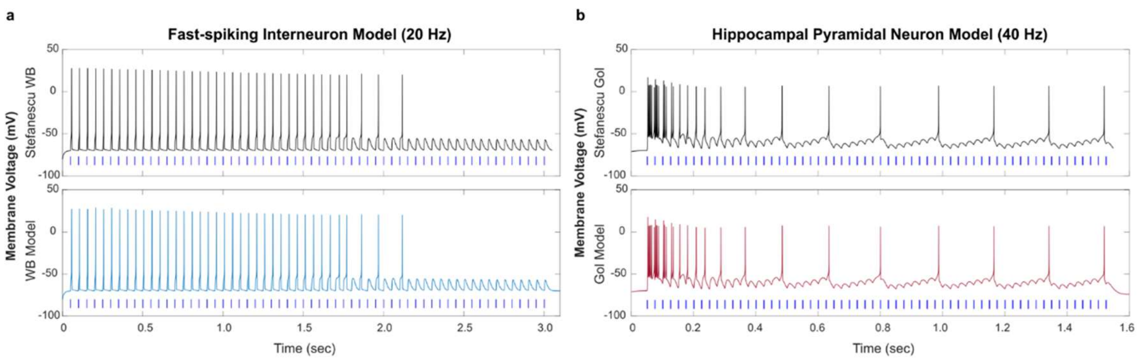


**Fig. S1.** Comparison of computational models. (a) Modeling ChR2wt spiking in a fast-spiking interneuron in response to 60, 2 ms pulses of optical stimulation at 20 Hz. Top panel: replicated modeling results from Fig. 6A, Stefanescu et al. (1). Bottom panel: results from our model with regular single-site stimulation. (b) Modeling ChR2wt spiking in a hippocampal pyramidal neuron in response to 60, 2 ms pulses of optical stimulation at 40 Hz. Top panel: replicated modeling results from Fig. 7A, Stefanescu et al. (1). Bottom panel: results from our model with regular single-site stimulation.

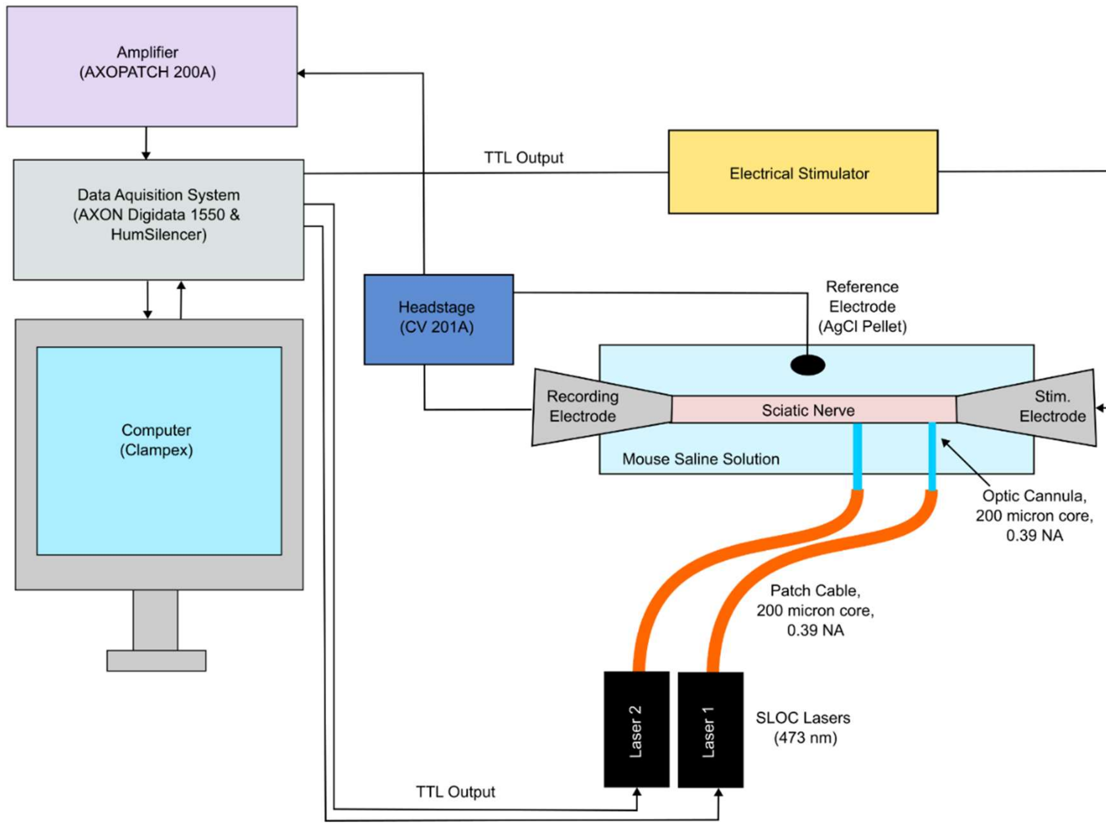


**Fig. S2.** Experimental setup for *ex vivo* compound action potential (CAP) recordings. Sciatic nerves were placed in a room temperature (~22°C) bath (P6/PH6, Warner Instruments) of pH buffered saline solution. Glass suction pipettes delivered electrical stimulation to the sciatic-side of the nerve and recorded compound action potentials (CAPs) from the tibial-side of the nerve. A battery-operated stimulator (ISO-FLEX) was used to deliver electrical stimulation via the suction electrode on the sciatic side of the nerve. Light pulses were delivered using two fiber optic cannulas (CFMC12L20, 0.39 NA, 200 µm core, Thorlabs) that were pressed flush against the nerve using two separate micromanipulators (MP-285, Sutter Instruments). Each cannula was connected to a 473 nm laser (BL473T8-100FC, Shanghai Laser Optics Century Co. (SLOC)) via a patch cable (M81L01, 0.39 NA, 200 µm core, Thorlabs) using a quick-release interconnect (ADAF2, Thorlabs). CAP recordings from electrical and optical stimulation were collected using a CV-201A headstage, an Axon Digidata 1550A, an Axopatch 200A amplifier with a gain of 0.5 and a lowpass Bessel filter at 5 kHz, an Axon Digidata 1550A digitizer, and Clampex Software (version 10.5.0.9, Molecular Devices). An Ag/AgCl pellet (E205, Warner Instruments) was submerged in the bath to complete the circuit and act as the reference electrode. All experiments were performed inside a faraday cage on an Axioskop 2 FS plus microscope (Zeiss). A long working distance air objective (A-Plan 2.5X/0.06, Zeiss) and camera (Part #BFLY-U3-23S6M-C, FLIR) were used to confirm the cannula positions. For each experiment, the cannulas were placed ~3 mm apart.

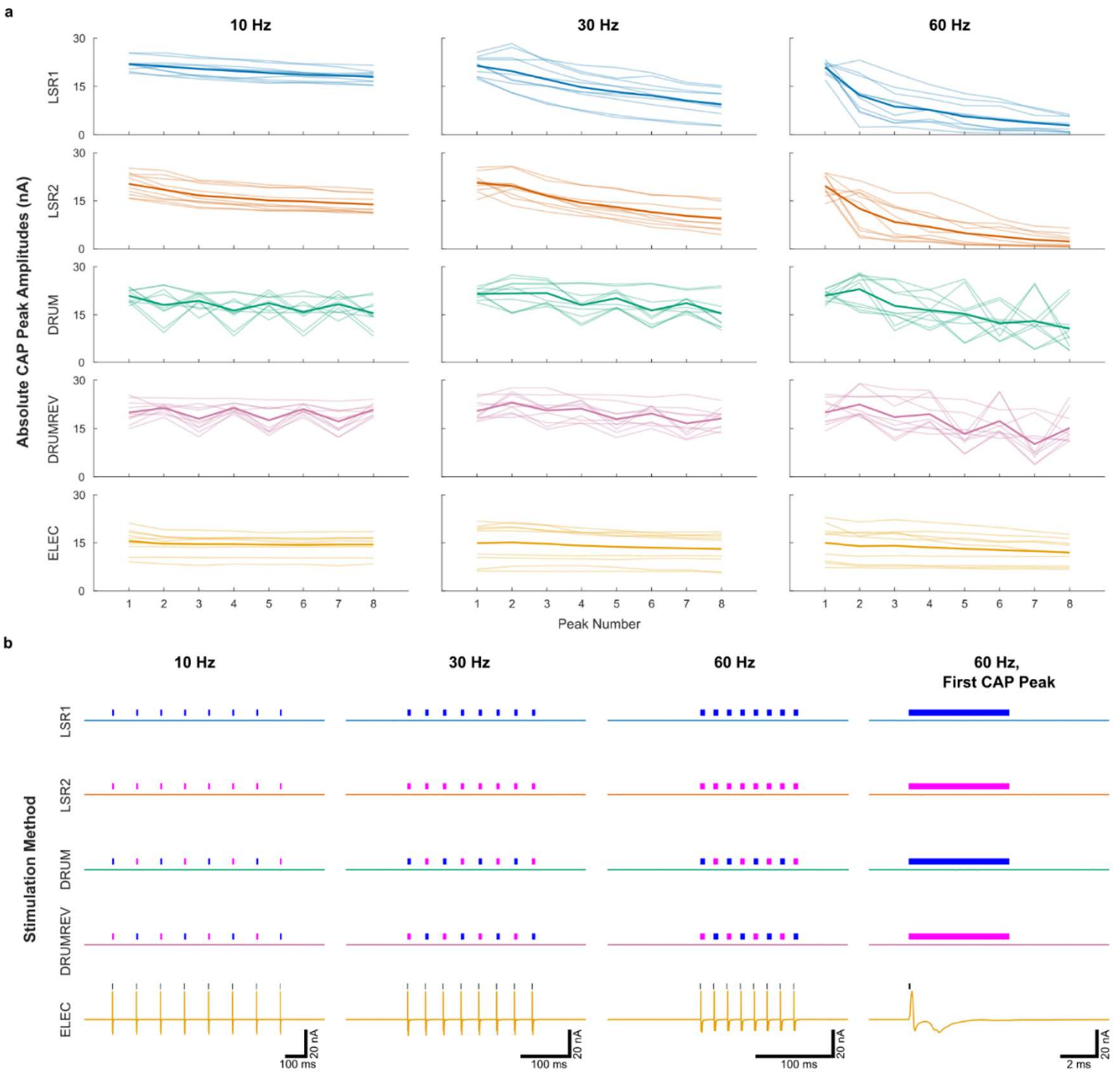


**Fig. S3.** CAP Peak Amplitudes from all experiments and CAP traces from a control experiment. (a) CAP Peak Amplitude traces as a function of peak number pooled by stimulation method and frequency. Lighter colored lines represent all individual measurements (taken from n = 5 mice for optical methods and n = 6 mice for electrical methods). The darker colored line represents the average of all the traces. (b) Representative *ex vivo* CAP traces from a control experiment. Recordings were taken at the maximum laser intensities used in the non-control experiments (LSR1 = 62 mW/mm$^2$, LSR2 = 379 mW/mm$^2$).

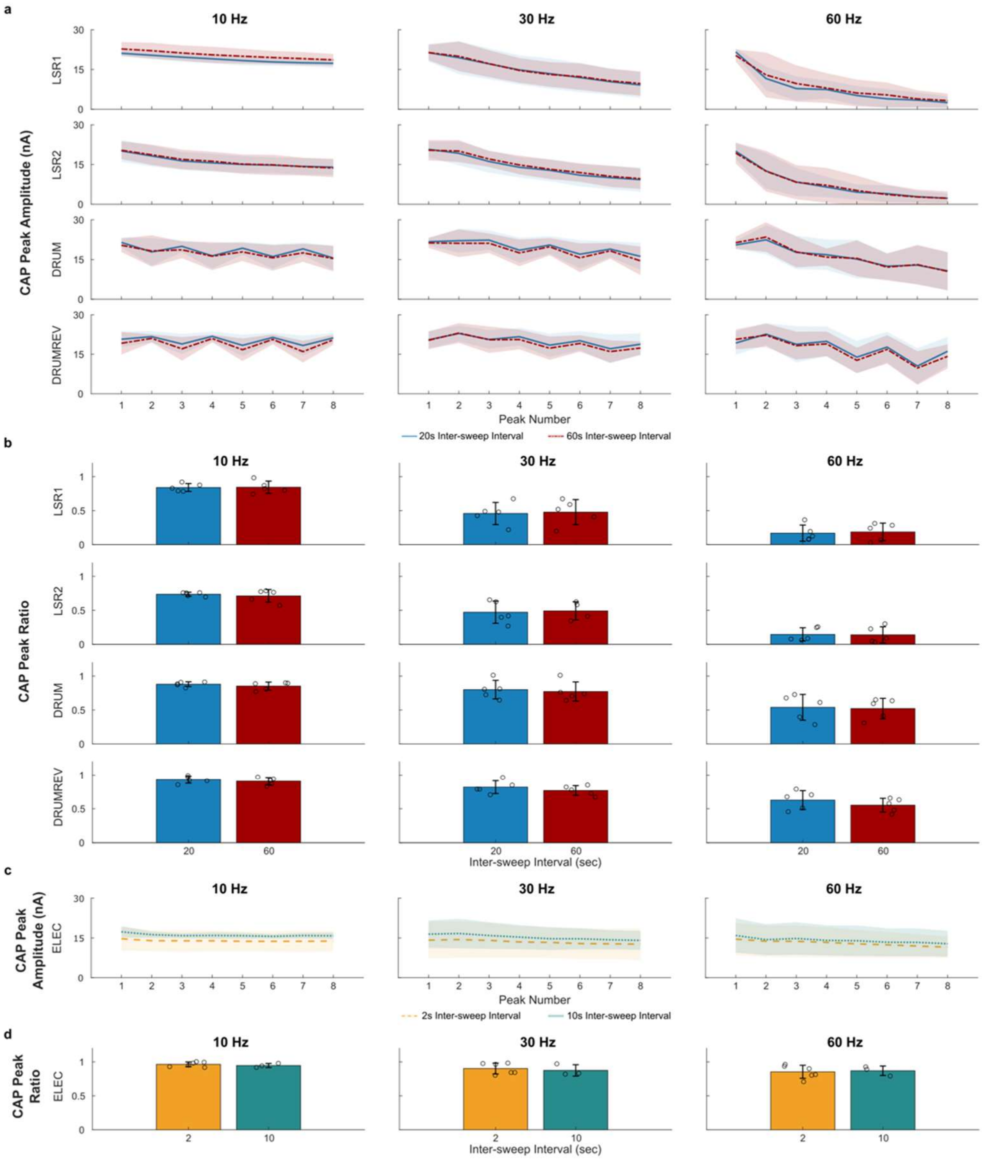


**Fig. S4.** Effect of inter-sweep interval on CAP peak amplitudes. (a) Average CAP Peak Amplitudes (pA) as a function of peak number for all optical stimulation methods. Solid and dot-dashed lines represent the average CAP peak amplitudes across eight successive stimulation pulses for the 20 and 60 second inter-sweep interval respectively. Transparent borders represent the standard deviation across multiple mice (n = 5 for all optical conditions except for LSR2, 30 Hz, and an inter-sweep interval of 60 seconds where n = 4). (b) CAP Peak Ratio for each optical inter-sweep interval. For each optical stimulation method and frequency, there were no statistically significant differences ($p > 0.05$) between the CAP Peak Ratios calculated at 20 and 60 second intervals. (c) Average CAP Peak Amplitudes (pA) as a function of peak number for electrical stimulation. Dashed and dotted lines represent the average CAP peak amplitudes across eight successive stimulation pulses for the 2 and 10 second inter-sweep interval respectively. Transparent borders represent the standard deviation across multiple mice (n = 6 for 2 second inter-sweep interval and n = 3 for the 10 second inter-sweep interval). (d) CAP Peak Ratio for each electrical inter-sweep interval. For each stimulation frequency, there were no statistically significant differences ($p > 0.05$) between the CAP Peak Ratios calculated at 2 and 10 seconds.

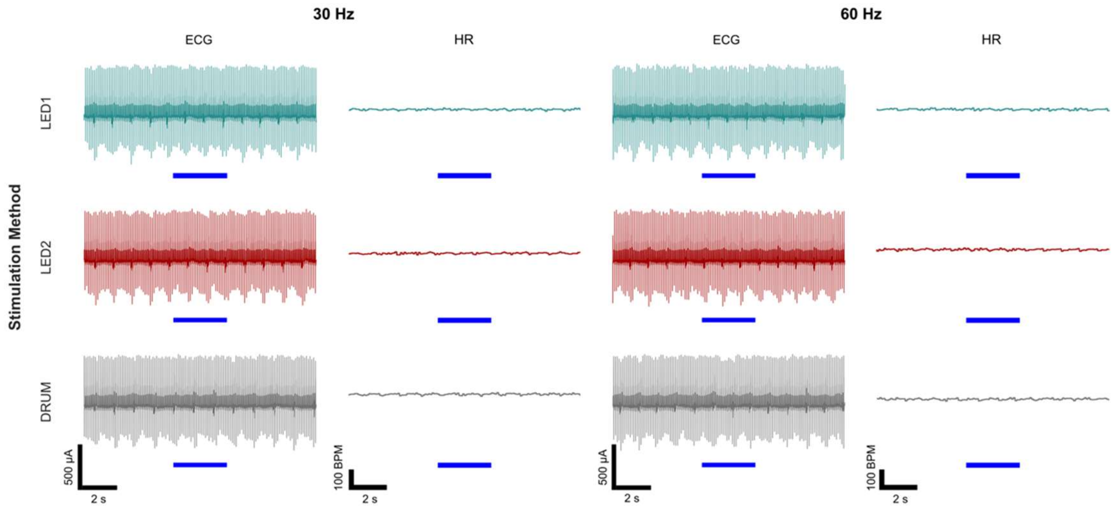


**Fig. S5.** *In vivo* optogenetic vagus nerve stimulation (VNS) in a control animal. Raw ECG traces and corresponding average heart rate (HR) during three seconds of optogenetic VNS in a mouse without ChR2 expression.

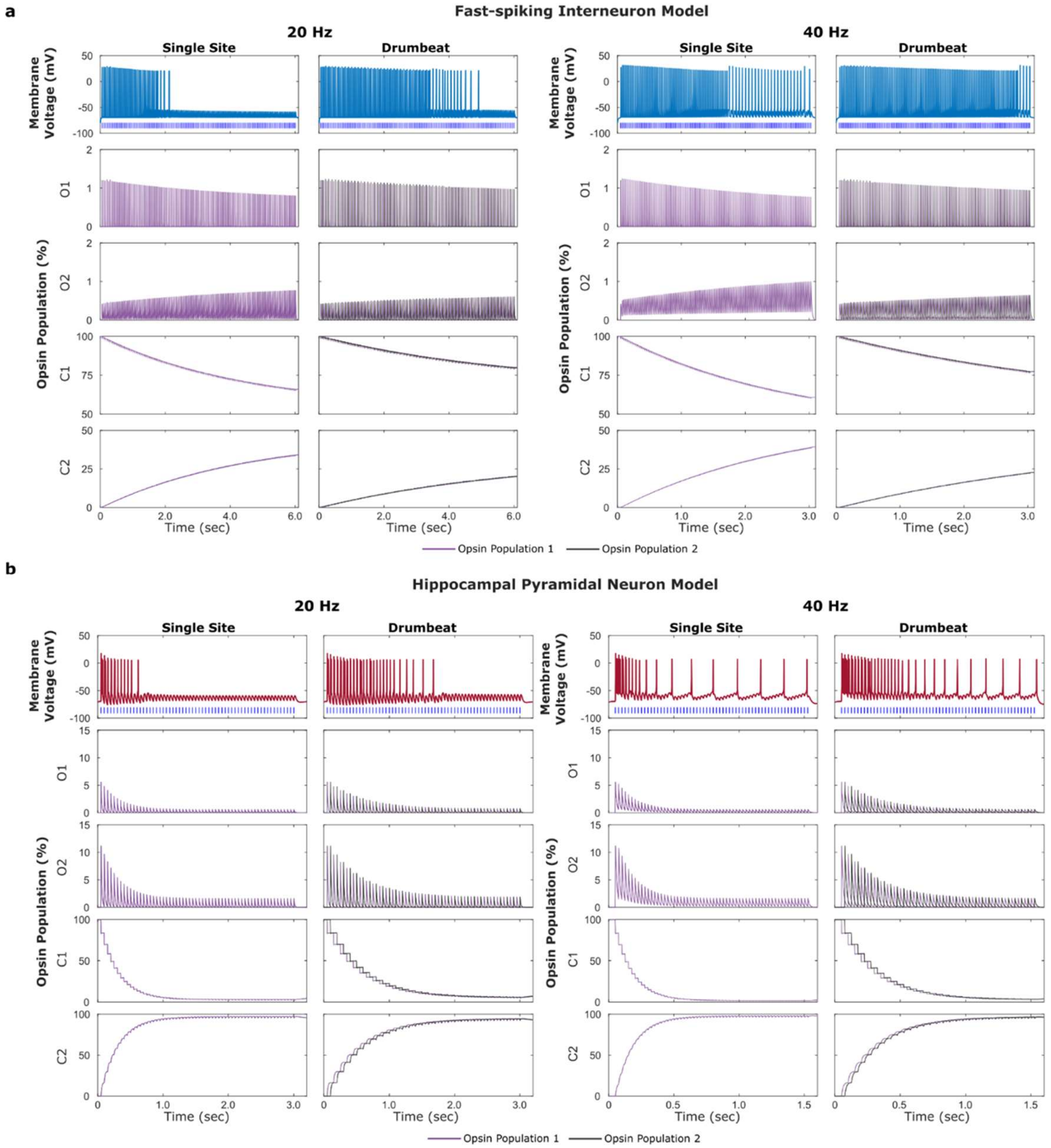


**Fig. S6.** Computational modeling of single-site and two-site drumbeat stimulation in fast-spiking interneurons and hippocampal pyramidal neurons. (a) Modeling ChR2wt spiking in a fast-spiking interneuron in response to 120, 2 ms pulses of single-site and drumbeat stimulation at 20 and 40 Hz. Lower panels show the fraction (%) of the total opsin population in each of the four states (O1, O2, C1, and C2) at a given time. (b) Modeling ChR2wt in a hippocampal pyramidal neuron in response to 60, 2 ms pulses of regular and drumbeat stimulation. Lower panels show the fraction (%) of the total opsin population in each of the four states (O1, O2, C1, and C2) at a given time.

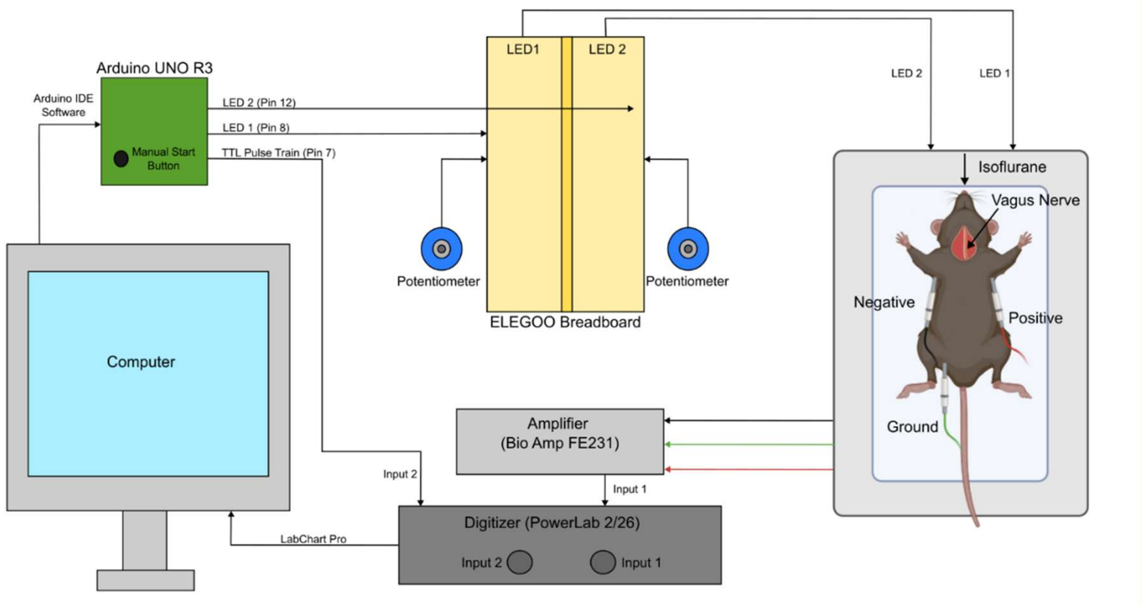


**Fig. S7.** Experimental setup for in-vivo electrocardiogram (ECG) recordings. For a given recording, the desired stimulation frequency (30 or 60 Hz) and stimulation method (LED1, LED2, or DRUM) was input into our custom Arduino IDE code (version 2.3.6, Arduino) and uploaded to an Arduino UNO R3. The UNO board was equipped with a manual button to trigger stimulation and three outputs: 1) TTL trigger output for LED1, 2) TTL trigger output for LED2, and 3) TTL trigger output for both LEDs which was routed to one of the digitizer's inputs. The TTL trigger for LED1 and LED2 was routed to an ELEGOO breadboard where the current input to each of the LEDs could be adjusted using a potentiometer (3590S-2-104L, Bourns). From the breadboard, the current was routed to the LEDs which were imbedded in our custom 3D-printed nerve cuffs (Fig. 4a). ECG readings were collected using leads placed in the mouse's armpits and groin. The ECG signal was then routed to a Bio Amp amplifier (FE231, AD Instruments) and a PowerLab2/26 digitizer (PL2602, AD Instruments). The output from the digitizer was collected using LabChart Pro (v8.1.30, AD Instruments). For each recording, we collected the raw ECG signal, the time averaged heart rate (beats per minute), and the TTL trigger for the LEDs. Mouse rendering was created in BioRender.
